# Unsupervised machine learning identifies latent pyrenoid states linked to mitotic remodeling defects and CO_2_-dependent growth

**DOI:** 10.64898/2026.08.25.746929

**Authors:** Koujiro Matsuo, Takashi Yamano

## Abstract

Biomolecular condensates that persist through cell division must be reorganized and inherited, yet it remains unclear whether subtle defects before division are associated with later organelle or growth phenotypes. We examined the *Chlamydomonas reinhardtii* pyrenoid, a liquid-like condensate that concentrates ribulose-1,5-bisphosphate carboxylase/oxygenase (Rubisco), the photosynthetic CO_2_-fixing enzyme. As part of the algal CO_2_-concentrating mechanism, the pyrenoid raises CO_2_ availability around Rubisco. We generated an RBCS1-mGold Rubisco reporter and developed an unsupervised image-analysis pipeline combining a convolutional autoencoder and a one-class support vector machine. Using 4,905 wild-type single-cell images, augmented 22-fold to 107,910 image instances, we defined the range of normal pyrenoid morphology. A combined machine-learning and visual screen of approximately 21,000 insertional mutants yielded 17 pyrenoid integrity mutants (*pim1*–*pim17*). Differential reconstruction-error maps highlighted local deviations from the wild-type reference, including phenotypes difficult to classify by eye. Four-dimensional live imaging showed defects in matrix dispersal, partitioning of Rubisco-containing foci, or pyrenoid recondensation in multiple *pim* strains. Growth assays identified broad defects and phenotypes that became more apparent as CO_2_ supply decreased. Insertion-site mapping nominated candidate loci, including *STT7*, which encodes a chloroplast kinase best known for regulating photosynthetic light harvesting. Independent *STT7*-edited lines lacked detectable STT7 accumulation and showed pyrenoid-region reconstruction-error patterns, supporting an association between impaired STT7 function and altered pyrenoid morphology. These findings show that unsupervised image screening can extend forward genetics to subtle pyrenoid phenotypes accompanied by mitotic remodeling or growth defects.

**Significance statement:** Membraneless organelles lack enclosing membranes, yet their organization must be maintained or rebuilt as cells divide or conditions change. We developed an unsupervised machine-learning screen for the *Chlamydomonas* pyrenoid, a phase-separated photosynthetic condensate that supports CO_2_ fixation. The screen recovered pyrenoid integrity mutants, including strains whose interphase pyrenoids appeared nearly normal but showed defects during mitotic remodeling or growth as CO_2_ supply decreased. Static images thus helped identify strains with subsequent mitotic or growth phenotypes, and insertion-site mapping nominated associated candidate loci. The approach extends forward genetics to dynamic cellular compartments whose informative phenotypes are difficult to define in advance.

## Introduction

Eukaryotic cells organize biochemical reactions not only within membrane-bound organelles but also within membraneless biomolecular condensates. Many such compartments form through multivalent macromolecular interactions and liquid-liquid phase separation, allowing selected proteins and nucleic acids to concentrate without enclosure by a lipid membrane (1–4). For condensates that persist through cell division, this mode of organization poses a distinctive cell-biological challenge. Their composition and material properties must be preserved or re-established as they undergo dissolution, intracellular redistribution, partitioning, and reassembly.

The pyrenoid of *Chlamydomonas reinhardtii* provides a powerful system for studying this problem in a physiologically important organelle. The pyrenoid concentrates ribulose-1,5-bisphosphate carboxylase/oxygenase (Rubisco), the enzyme that catalyzes photosynthetic CO_2_ fixation. As a central component of the algal CO_2_-concentrating mechanism (CCM), it increases CO_2_ availability around Rubisco and thereby supports efficient carbon fixation (5, 6). In *Chlamydomonas*, the pyrenoid matrix forms through multivalent interactions between Rubisco and the intrinsically disordered linker EPYC1 and behaves as a liquid-like condensate that reorganizes during cell division (5–8). Additional Rubisco-binding and matrix-associated proteins connect the matrix to surrounding subcompartments, including the starch sheath and matrix-traversing thylakoid membranes (9–13).

These studies have established core principles of pyrenoid assembly, but less is known about how pyrenoid organization is maintained during cell-cycle progression and changes in CO_2_ availability. Recent work showing that the kinase KEY1 tunes pyrenoid condensate size and cell-cycle-dependent dissolution supports the view that pyrenoid remodeling is actively regulated rather than a passive consequence of matrix assembly (14). Defects in these processes need not cause complete pyrenoid loss. They may instead appear as small changes in matrix shape, position, intensity, or spatial organization during interphase and become evident only when the pyrenoid disperses and reassembles during mitosis or functions under reduced CO₂ supply.

Forward genetic screens based on growth or visual morphology are well suited to isolating mutants with obvious physiological defects or gross pyrenoid abnormalities. They are less suited to phenotypes that are difficult to define in advance, particularly when an abnormality is distributed across several image features or becomes apparent only at a later cell-cycle stage or under reduced CO_2_. This problem is especially relevant to phase-separated organelles, in which modest changes in interaction strength, composition, or spatial organization may be tolerated under one condition but become consequential during dynamic remodeling.

Image-based machine learning can complement manual screening by identifying cells that deviate from a learned range of normal morphology without requiring the informative feature to be specified beforehand. Unsupervised novelty-detection and autoencoder-based approaches have been used to detect rare or unexpected cellular states in high-content microscopy and medical imaging (15, 16). Here, we combined a convolutional autoencoder with a one-class support vector machine to define the range of wild-type pyrenoid morphology in a high-dimensional image representation (17). We used this pipeline together with visual inspection to screen insertional mutants and asked whether subtle features in static interphase images were associated with pyrenoid remodeling during mitosis and with CO_2_-dependent growth.

## Results

### Development of an unsupervised anomaly-detection pipeline

The *Chlamydomonas* pyrenoid is visible in bright-field images and, at the ultrastructural level, comprises a Rubisco-rich matrix surrounded by a starch sheath and traversed by pyrenoid tubules that are continuous with the thylakoid membrane system (Fig. 1A). To identify pyrenoid phenotypes that are difficult to define by visual criteria, we generated the RG2 reporter strain, which expresses a Rubisco small subunit 1 (RBCS1)-mGold fusion protein in a wild-type (WT) CC-125 background. StarDist2D segmentation followed by mask quality filtering yielded 4,905 WT single-cell images for defining the reference distribution of pyrenoid morphology (Fig. 1B) (18).

**Figure 1.**
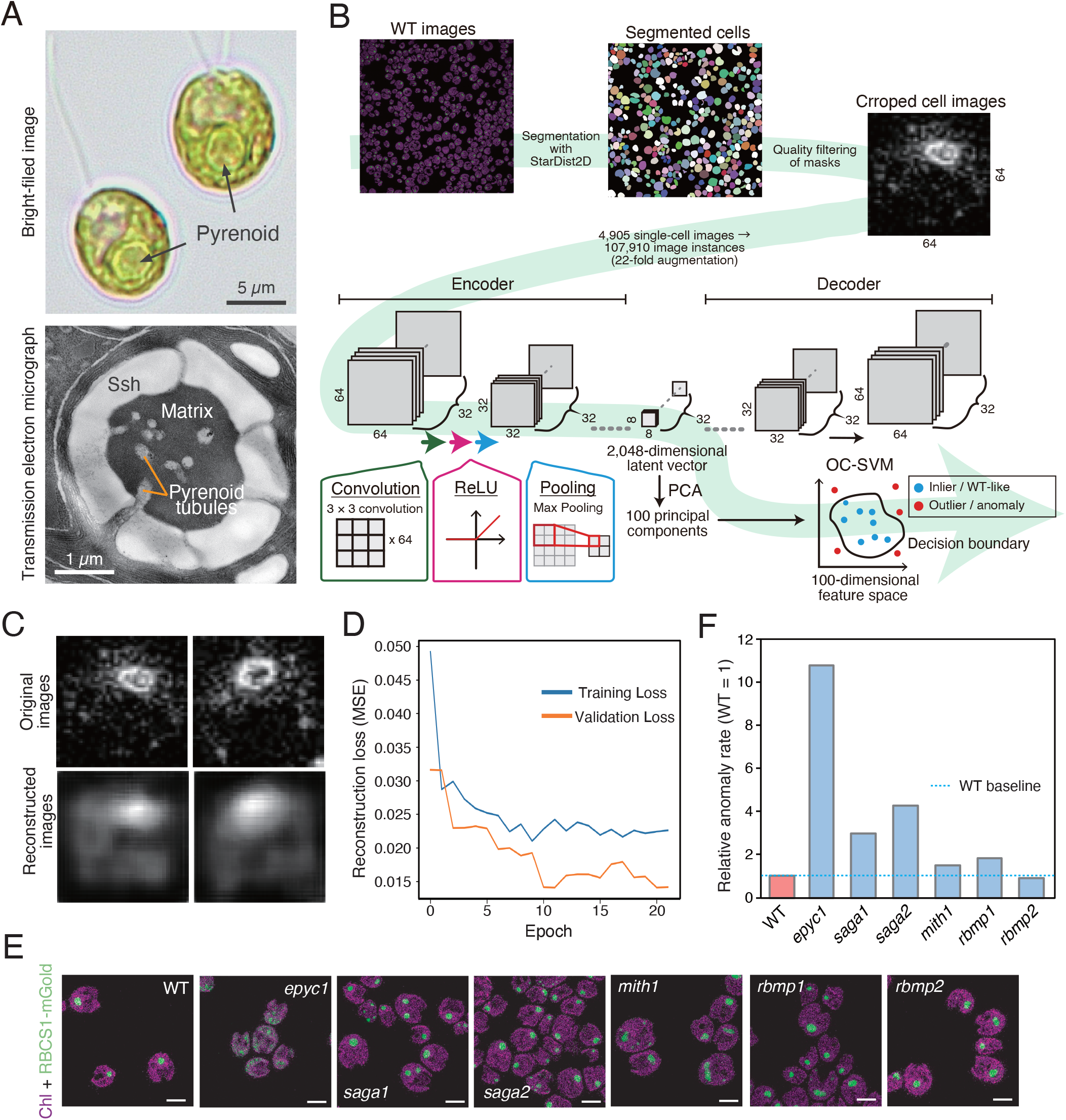
Unsupervised anomaly detection of pyrenoid morphology. **(A)** Representative bright-field image of wild-type (WT) *Chlamydomonas reinhardtii* cells (upper) and transmission electron micrograph of a pyrenoid cross-section (lower), showing the Rubisco-rich matrix, starch sheath (Ssh), and matrix-traversing pyrenoid tubules. Scale bars, 5 µm (upper) and 1 µm (lower). **(B)** Overview of the convolutional autoencoder–one-class support vector machine (CAE–OC-SVM) pipeline. A total of 4,905 quality-filtered WT single-cell images yielded 107,910 image instances after 22-fold augmentation. Images were encoded into a 2,048-dimensional latent vector, reduced to 100 principal components, and classified relative to the estimated WT support. **(C)** Representative WT model-input images (upper) and CAE reconstructions (lower). **(D)** Training and validation mean squared reconstruction loss; training stopped after 22 epochs. **(E)** Representative images of WT and established pyrenoid-related mutants used for biological calibration. Magenta, chlorophyll autofluorescence; green, RBCS1-mGold. Scale bars, 5 µm. **(F)** Anomaly rates relative to WT (WT = 1.0); the dashed line indicates the WT-relative baseline.

We built an unsupervised pipeline consisting of a convolutional autoencoder (CAE) followed by one-class support vector machine (OC-SVM) classification (Fig. 1B) (15–17). Source image files were partitioned before augmentation so that images from the same field could not enter both training and validation partitions. Twenty-two-fold augmentation within the assigned partitions expanded the 4,905 source images to 107,910 image instances. The CAE encoded each normalized 64 × 64-pixel image into a 2,048-dimensional latent vector. Principal component analysis (PCA) reduced this representation to 100 principal components, and the OC-SVM estimated the support of WT morphology in the reduced feature space.

This architecture represented each cell through learned image features rather than a single predefined morphological descriptor. Representative CAE outputs retained the dominant pyrenoid-centered intensity pattern while smoothing fine spatial detail (Fig. 1C). Training and validation mean squared reconstruction losses decreased and stabilized, and the early-stopping criterion terminated training after 22 epochs (Fig. 1D). The OC-SVM parameter *ν* was set to 0.05. Because *ν* constrains the fraction of training observations permitted outside the estimated WT support, the approximately 5% WT anomaly rate was treated as a designed baseline rather than an independent measure of classifier accuracy (17).

We then applied the trained pipeline to a panel of established pyrenoid-related mutants as a biological calibration (Fig. 1E and F) (5, 9–12). Relative anomaly rate was calculated by dividing each strain’s anomaly rate by the WT anomaly rate. The resulting values were approximately 10.8 for *epyc1*, 3.0 for *saga1*, 4.3 for *saga2*, 1.5 for *mith1*, 1.8 for *rbmp1*, and 0.9 for *rbmp2*, compared with 1.0 for WT. The strong response of *epyc1* was consistent with the established requirement for EPYC1 in matrix formation (5–8), whereas the other strains spanned intermediate to near-baseline values. Thus, the pipeline responded to a range of established pyrenoid phenotypes. Because these mutants were used for biological calibration rather than as a held-out test set, the panel was not used to estimate sensitivity or specificity.

### A combined machine-learning and visual screen identifies pyrenoid integrity mutants

We applied the CAE-OC-SVM pipeline to a random insertional mutant collection using a hybrid strategy that combined anomaly scoring with visual inspection. Approximately 21,000 mutants were first surveyed by microscopy to exclude strains with obvious growth defects or complete loss of the pyrenoid reporter, thereby enriching for colonies with ambiguous or subtle phenotypes. This prescreen left 1,728 candidate strains for image-based analysis, which were grouped 24 colonies per well to generate 72 pools for the primary screen (Fig. 2A).

**Figure 2.**
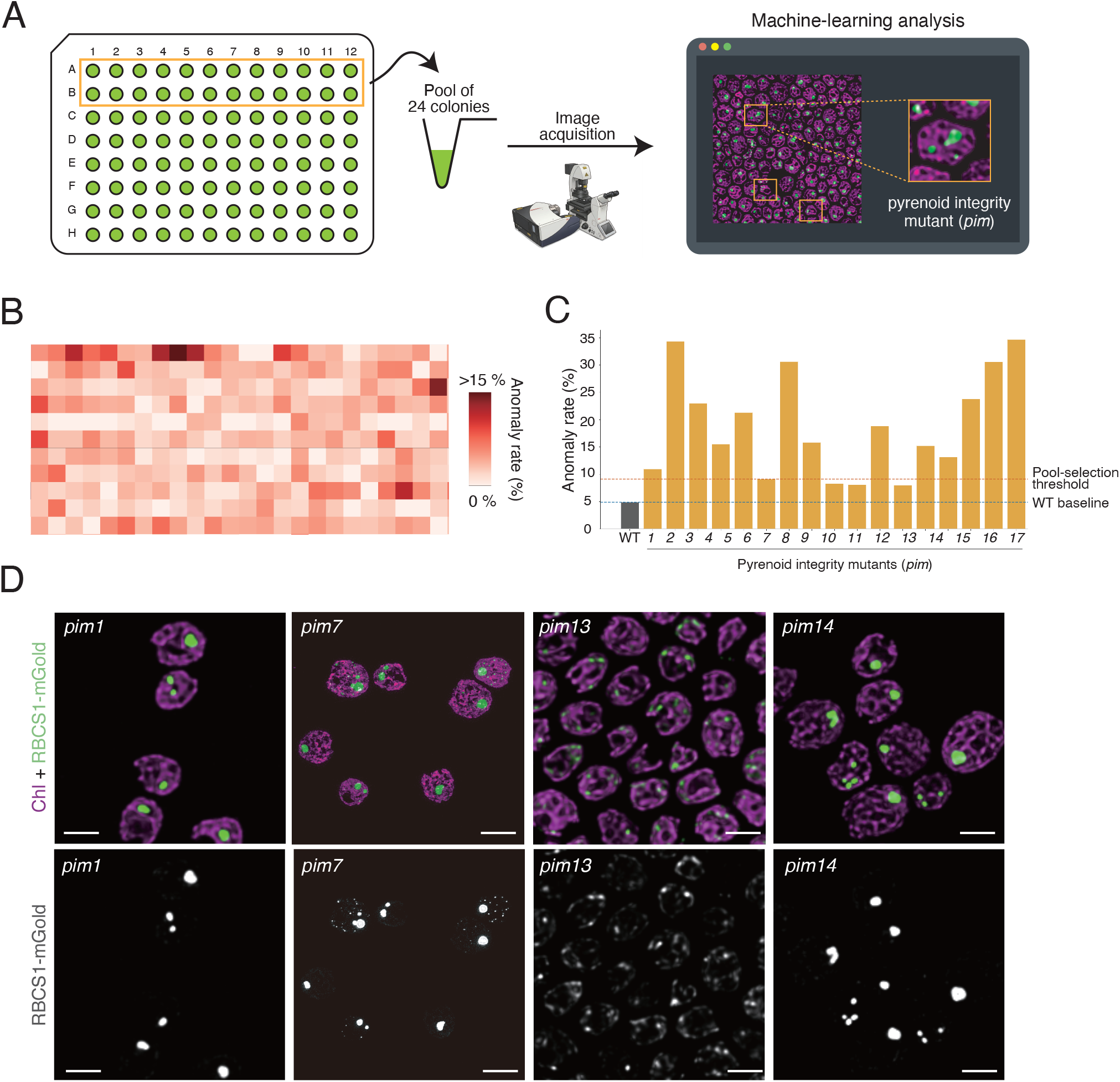
Combined machine-learning and visual screen for pyrenoid integrity mutants. **(A)** Screening workflow. An initial visual prescreen of approximately 21,000 insertional mutants yielded 1,728 candidate strains, which were grouped 24 colonies per well to generate 72 pools for CAE–OC-SVM analysis. **(B)** Heat map of anomaly rates for the 72 pools. Pools exceeding the matched WT baseline by approximately 4.2 percentage points were selected for secondary analysis. **(C)** Anomaly rates of the 17 recovered pyrenoid integrity mutants (*pim1*–*pim17*). The blue dashed line indicates the WT baseline (approximately 4.9%), and the orange dashed line indicates the primary pool-selection threshold. Visual confirmation also contributed to final strain selection. **(D)** Representative confocal images of *pim1*, *pim7*, *pim13*, and *pim14*. Top, merged chlorophyll autofluorescence (magenta) and RBCS1-mGold (green); bottom, RBCS1-mGold alone in grayscale. Scale bars, 5 µm.

Pooled samples were imaged and their anomaly rates were summarized as a heat map (Fig. 2B). Pools whose anomaly rates exceeded the matched WT baseline by approximately 4.2 percentage points were taken forward for single-colony analysis. Individual colonies from selected pools were then re-imaged. A strain was retained when its anomaly rate reproducibly exceeded the WT baseline by the predefined cutoff in repeated assays or when repeated visual inspection revealed a consistent pyrenoid phenotype (Fig. 2C). The second route was important for phenotypes dominated by a simple feature that was not strongly emphasized by the CAE score. For example, *pim8* showed only a modest anomaly rate but a reproducibly small pyrenoid.

The combined screen yielded 17 pyrenoid integrity mutants, designated *pim1*–*pim17* (Fig. 2C and D). Their anomaly rates spanned a broad range, from near-threshold values for some strains to >30% for *pim2* and *pim17*, whereas the WT baseline was approximately 4.9% (Fig. 2C). Representative examples included small or multiple pyrenoid foci (*pim1*), diffuse or dispersed matrix-associated signal (*pim7*), severe matrix disruption (*pim13*), and enlarged or displaced foci (*pim14*) (Fig. 2D). Because the workflow included pooling, a visual prescreen, and a secondary confirmation step, it was designed to enrich reproducible pyrenoid phenotypes rather than estimate the frequency of all possible defects in the original library.

### Differential reconstruction-error mapping highlights spatially localized image deviations

The OC-SVM score identifies cells outside the estimated WT support but does not indicate which image regions differ from the WT reference. To localize such differences, we calculated for each cell a raw reconstruction-error map as the absolute pixel-wise difference between the input image and its CAE reconstruction (Fig. 3A and B) (16). WT images were divided into an 80% reference subset and a separate 20% comparison subset. The mean raw reconstruction-error map calculated from the 80% reference subset was used as the WT reference map. Subtracting this reference from the mean raw error map of the 20% comparison subset produced the mean WT differential reconstruction-error map shown in Fig. 3C. Its near-zero spatial pattern indicates that the average reconstruction-error distribution in the comparison subset closely matched that of the reference subset.

**Figure 3.**
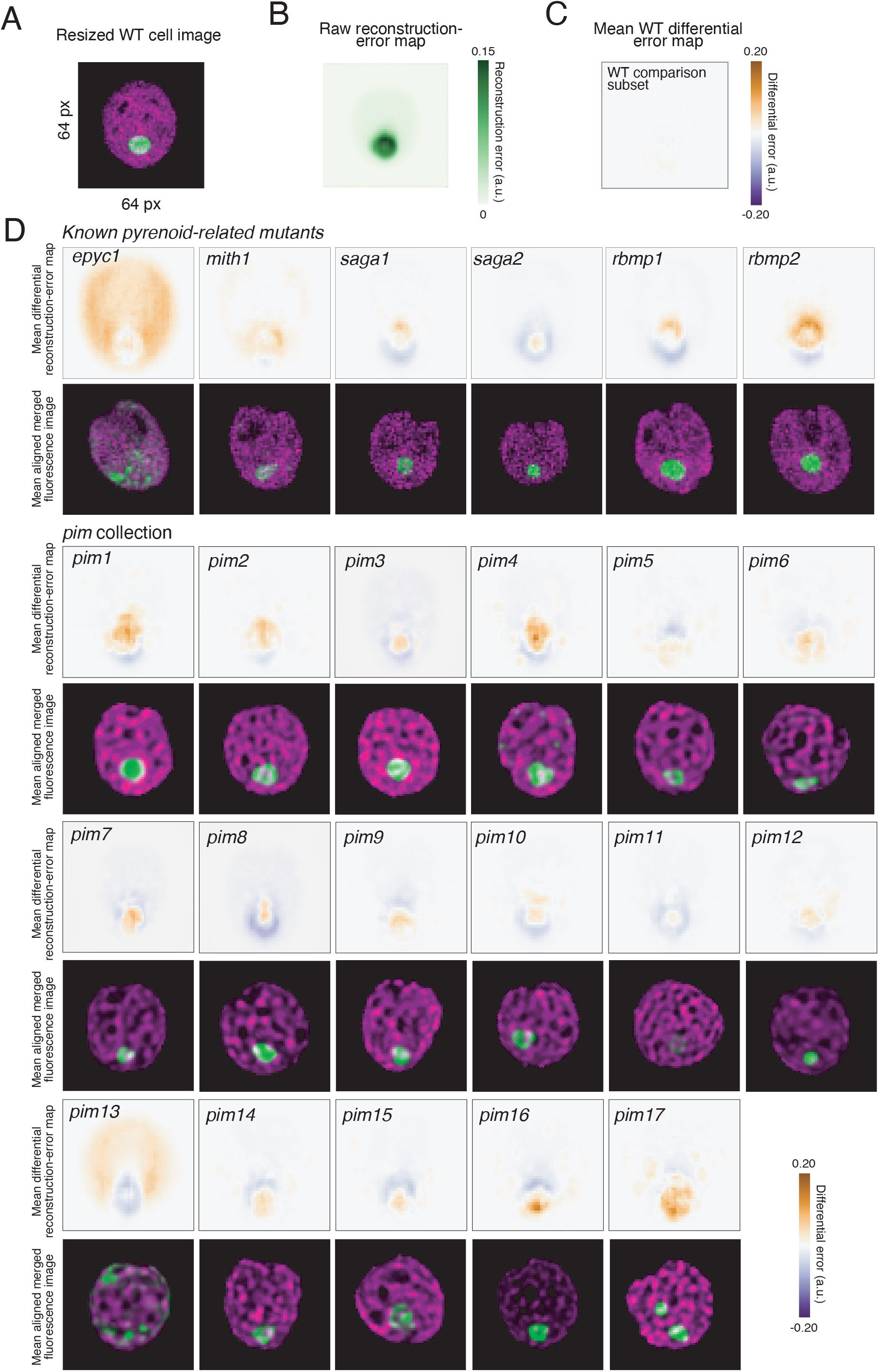
Differential reconstruction-error mapping highlights spatially localized image deviations. **(A)** Representative resized WT fluorescence image. **(B)** Raw reconstruction-error map, calculated as the absolute pixel-wise difference between the model-input image and its CAE reconstruction. **(C)** Mean WT differential reconstruction-error map generated by subtracting the mean raw error map of an 80% WT reference subset from that of a separate 20% WT comparison subset. The near-zero pattern indicates similar average spatial error distributions in the two subsets. **(D)** Mean differential reconstruction-error maps and corresponding mean merged fluorescence images for established pyrenoid-related mutants and *pim1*–*pim17*. Chlorophyll autofluorescence is shown in magenta and RBCS1-mGold in green. All differential maps use the same −0.20 to 0.20 a.u. scale. Positive and negative values indicate reconstruction error above or below the WT reference and are not assigned direct structural or molecular meaning. Mean fluorescence images are shown for morphological context.

Differential maps for known mutants and *pim* strains were calculated using the same WT reference map. Values near zero indicate locations at which reconstruction error is similar to the WT reference. Positive values indicate locations with higher reconstruction error than the WT reference, whereas negative values indicate locations with lower reconstruction error. The sign of a value does not by itself indicate the gain, loss, or displacement of a particular structure, and the maps are not causal saliency maps. Rather, they identify image regions that the WT-trained CAE reconstructs differently from the WT reference.

To summarize recurring genotype-level patterns, we calculated a mean differential reconstruction-error map and a mean merged fluorescence image from the corresponding set of aligned single-cell images for each strain (Fig. 3D). The mean fluorescence images provide morphological context for the spatial error patterns. Among the known pyrenoid-related mutants, epyc1 showed the broadest positive differential-error pattern, whereas *mith1*, *saga1*, *saga2*, *rbmp1*, and *rbmp2* showed weaker or more localized patterns. Applying the same analysis to the *pim* collection revealed local deviations even in strains whose fluorescence images were difficult to distinguish from WT by eye. For example, *pim1*, *pim3*, and *pim5* showed compact positive regions near the pyrenoid, *pim7* showed a displaced pattern, and *pim13* displayed broad deviations extending beyond the central matrix signal. These maps provide spatial hypotheses for follow-up analysis but do not establish the molecular identity or physical basis of the underlying defect.

### Pyrenoid remodeling and growth phenotypes across the *pim* collection

We next examined pyrenoid behavior during cell division by four-dimensional live imaging of RBCS1-mGold. Cells were immobilized in low-melting-point agarose and followed by confocal laser-scanning microscopy (Fig. 4A). Selected frames are shown for WT and all 17 *pim* strains. In WT cells, the matrix dispersed into numerous Rubisco-containing foci during the first division, redistributed through the second division, and recondensed into a dominant pyrenoid in each daughter lineage, consistent with the dynamic behavior reported previously (6). Multiple *pim* strains showed incomplete matrix dispersal, persistence of large foci, altered numbers or spatial distributions of Rubisco-containing foci, or unequal partitioning of the reporter signal. Because Fig. 4A presents selected stages, these sequences document representative remodeling phenotypes rather than providing a quantitative comparison among strains.

**Figure 4.**
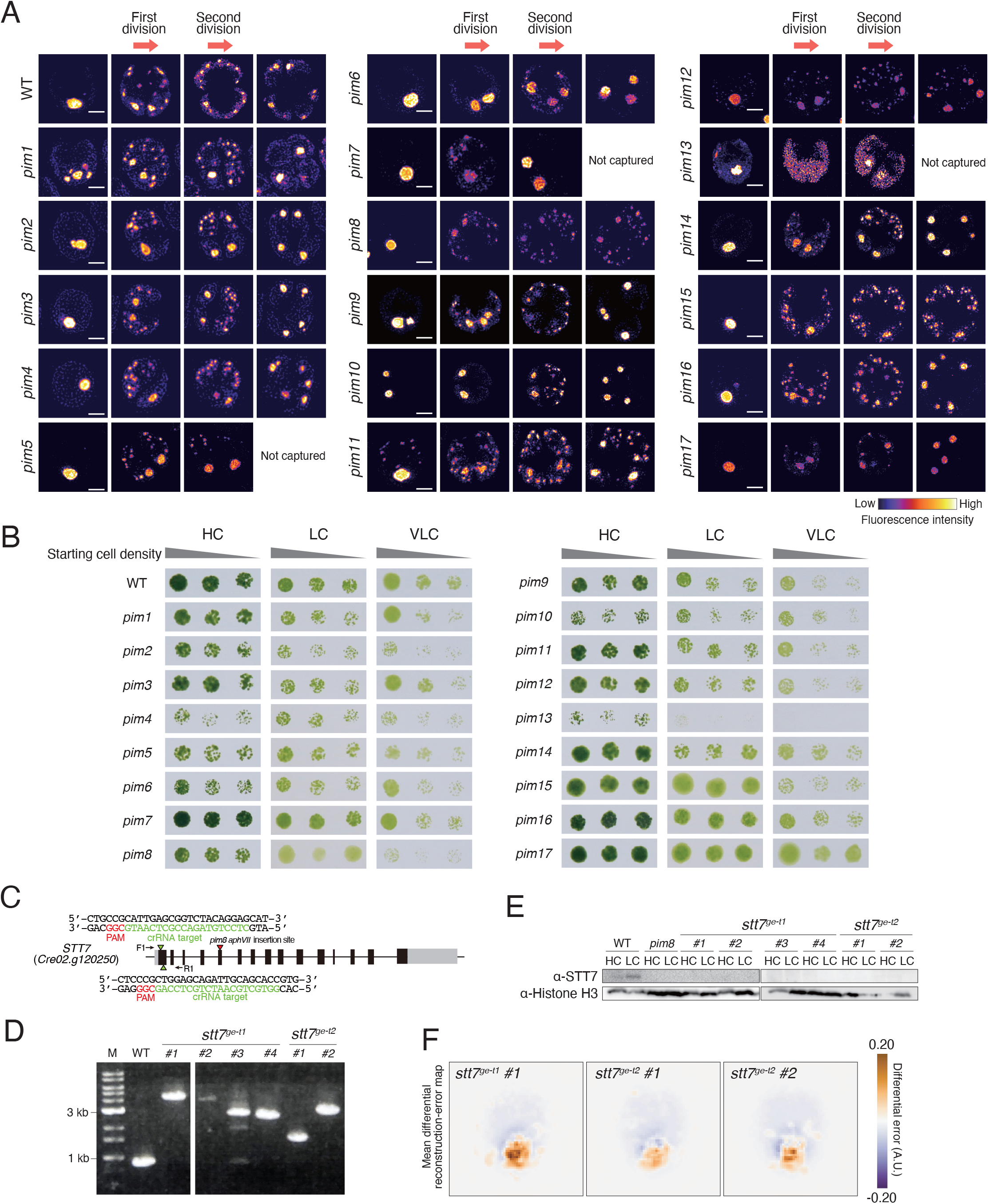
Mitotic and growth phenotypes of *pim* strains and analysis of the STT7 candidate locus. **(A)** Selected frames from four-dimensional imaging of RBCS1-mGold during cell division in WT and *pim1*–*pim17*, arranged from left to right in temporal order. Frames were selected to represent comparable stages rather than identical elapsed times across strains. Red arrows indicate the approximate first and second division intervals. RBCS1-mGold fluorescence is displayed using the indicated pseudocolor scale. “Not captured” indicates that a corresponding stage was not available in the recorded time series. Scale bars, 5 µm. **(B)** Spot assays of WT and *pim1*–*pim17* under high-CO_2_ (HC), low-CO_2_ (LC), and very-low-CO_2_ (VLC) regimes. Within each condition, inoculum densities decrease from left to right (OD_730_ = 0.15, 0.08, and 0.03). **(C)** *STT7* gene model (*Cre02.g120250*) showing the mapped *aphVII* insertion in *pim8*, two crRNA target sites, and F1/R1 primer positions. PAM and crRNA target sequences are shown in red and green, respectively. **(D)** PCR genotyping of WT and independent clones edited at the first (*stt7*^ge-t1^, clones #1–#4) or second (*stt7*^ge-t2^, clones #1 and #2) target site. M, DNA size marker; white gaps mark boundaries between separately displayed gel sections. **(E)** Immunoblot analysis of STT7 in WT, *pim8*, and the six independent edited clones under HC and LC. Histone H3 was used as a loading control; boundaries between separately displayed blot sections are indicated. **(F)** Mean differential reconstruction-error maps for three selected *STT7*-edited clones, calculated with the WT reference used in Fig. 3 and displayed on the same −0.20 to 0.20 a.u. scale. Positive and negative values are not assigned direct structural or molecular meaning.

To determine whether the image and division phenotypes were accompanied by growth differences, we performed spot assays on MOPS-P medium under operationally defined high-CO_2_ (HC), low-CO_2_ (LC), and very-low-CO_2_ (VLC) regimes. Three inoculum densities were tested for each strain (OD_730_ = 0.15, 0.08, and 0.03; Fig. 4B). The collection did not show a uniform CO_2_-specific response. *pim4*, *pim5*, and *pim13* grew more slowly than WT even under HC. Under LC, *pim13*, which carried an insertion in *EPYC1* and lacked a normal matrix, showed the strongest defect, whereas *pim4* and *pim5* remained comparatively weak. Under VLC, reduced growth was also evident for *pim2*, *pim4*, *pim5*, *pim6*, *pim8*, *pim9*, *pim10*, *pim11*, *pim12*, *pim13*, and *pim15*. The reduction was most apparent under VLC for *pim2*, *pim6*, *pim8*, *pim9*, *pim10*, *pim11*, *pim12*, and *pim15*, whereas the phenotypes of *pim4*, *pim5*, and *pim13* extended from HC to the lower-CO_2_ regimes. *pim1* and *pim3* showed mild or intermediate changes, and *pim7*, *pim14*, *pim16*, and *pim17* remained close to WT across the tested conditions. Because MOPS-P contains no added organic carbon source, these assays assessed photoautotrophic growth across the imposed CO_2_ regimes. They do not, however, distinguish a specific defect in the CCM from broader effects on photosynthesis, chloroplast physiology, or cellular growth.

### Insertion-site mapping nominates candidate loci

Whole-genome sequencing mapped one or more insertions associated with each *pim* strain (Table 1). Except where independent genetic evidence was available, the mapped loci were treated as candidates rather than causal genes. *pim13* carried an insertion in *EPYC1* (*Cre10.g436550*), consistent with its severe matrix phenotype and the established role of EPYC1 in pyrenoid assembly (5–8). Single annotated candidate loci were associated with *pim2* (a PHD-finger domain protein), *pim3* (P4H7), *pim5* (SABC1), *pim7* (OSB2), *pim9* (a putative transferase), *pim10* (SDR14/SDR7C), *pim11* (SRP19), and *pim15* (GT90F32). *pim1* (*Cre11.g481115*) and *pim6* (*Cre06.g283700*) were associated with loci encoding proteins of unknown function, whereas *pim4*, *pim12*, *pim16*, and *pim17* carried intergenic insertions. Multiple mapped insertions were present in *pim8*, at *STT7/CDPK7* (*Cre02.g120250*), *CAV8* (*Cre12.g532350*), and *TIM13* (*Cre16.g650800*), and in *pim14*, at *NSG11* (*Cre07.g339050*) and a uridine kinase locus (*Cre13.g573800*). These mappings nominate loci for further analysis but do not establish the molecular basis of the corresponding phenotypes.

**Table 1.** Mapped insertion sites and candidate loci associated with *pim* strains.

| Strain | Locus ID (Phytozome v5.6) | Gene Name / Annotation | Previously reported CCM-interactome associations (bait; WD score) |
| --- | --- | --- | --- |
| *pim1* | *Cre11.g481115* | Hypothetical protein | — |
| *pim2* | *Cre08.g380151* | PHD-finger domain protein | — |
| *pim3* | *Cre14.g626200* | Prolyl 4-hydroxylase 7 (P4H7) | — |
| *pim4* | Intergenic | — | — |
| *pim5* | *Cre06.g273750* | SABC1 (Chloroplast sulfate transporter) | NAR1.2 (25.2), LCIA (2.9), CAH3 (2.9) |
| *pim6* | *Cre06.g283700* | Hypothetical protein (OG0015123) | — |
| *pim7* | *Cre12.g483720* | OSB2 (Organelle-specific ssDNA-binding protein 2) | — |
| *pim8* | *Cre02.g120250* | STT7 / CDPK7 (Ser/Thr protein kinase) | CAH3 (208.6), RCA1 (8.5), PSAH (6.0) |
| *pim8* | *Cre12.g532350* | CAV8 (Ion transporter / Polycystin) | PSAH, LCI11, EPYC1, SBE3, LCI24, LCIC, CAH6, PSAK, HST1, PSBP4 |
| *pim8* | *Cre16.g650800* | TIM13 (Mitochondrial translocase subunit) | CCP1 (7.5) |
| *pim9* | *Cre06.g272750* | Putative transferase | — |
| *pim10* | *Cre08.g380550* | SDR14 / SDR7C (Short-chain dehydrogenase/reductase) | — |
| *pim11* | *Cre02.g095070* | SRP19 (Signal recognition particle subunit) | — |
| *pim12* | Intergenic | — | — |
| *pim13* | *Cre10.g436550* | EPYC1 / LCI5 (Pyrenoid component) | — |
| *pim14* | *Cre07.g339050* | NSG11 (Actin-depolymerizing factor homology domain) | — |
| *pim14* | *Cre13.g573800* | Uridine kinase | — |
| *pim15* | *Cre17.g716350* | GT90F32 (Glycosyltransferase family 90) | — |
| *pim16* | Intergenic | — | — |
| *pim17* | Intergenic | — | — |

Locus IDs refer to *Chlamydomonas reinhardtii* genome assembly v5.6 (Phytozome). "Intergenic" indicates that the insertion site does not overlap with any annotated gene. The Interaction column lists bait proteins for which the corresponding gene product was identified as a putative interactor by affinity-purification mass spectrometry, with WD-scores (24) shown in parentheses where available. Em dashes (—) indicate not determined or not applicable.

### Independent *STT7*-edited lines support *STT7* as a candidate locus associated with *pim8*

Because *pim8* contained three mapped insertions, the insertion in *STT7* was not considered causal on the basis of mapping alone. We examined *STT7* further using independent lines edited at two crRNA target sites in the *STT7* locus (Fig. 4C). PCR genotyping identified six independent edited clones, four from the first target and two from the second target, with amplicon sizes that differed from WT (Fig. 4D). Immunoblotting detected STT7 in WT under both HC and LC but not in *pim8* or any of the six edited clones under the conditions tested (Fig. 4E). Mean differential reconstruction-error maps from three selected edited clones showed localized deviations in the pyrenoid region when calculated using the same WT reference as in Fig. 3 (Fig. 4F). The independent lesions, loss of detectable STT7 accumulation, and shared image-level phenotype support an association between STT7 loss and altered pyrenoid morphology. They do not establish that the *STT7* lesion is the sole cause of the *pim8* phenotype or that STT7 acts directly on the pyrenoid. Effects mediated through photosynthetic electron transport, thylakoid organization, or broader chloroplast physiology remain possible (19, 20).

## Discussion

This study presents a combined machine-learning and visual strategy for finding subtle pyrenoid phenotypes in a large insertional mutant collection. The CAE-OC-SVM pipeline represented WT morphology in a high-dimensional image space, and the 17 recovered *pim* strains collectively displayed interphase, mitotic, and growth phenotypes. An important biological observation is that several strains with subtle interphase image phenotypes also showed abnormal pyrenoid remodeling in representative division-stage time series. Growth assays further identified both broad growth defects and phenotypes that became more apparent as CO_2_ supply decreased.

We use the term “latent pyrenoid state” operationally to describe a subtle or nearly normal interphase image phenotype observed in a strain that also shows a remodeling or growth phenotype. The term does not imply that we directly measured a material property or a causal state. One possible interpretation is that some image patterns reflect partial changes in matrix organization. The *Chlamydomonas* matrix is assembled through multivalent Rubisco–EPYC1 interactions and behaves as a liquid-like condensate (5–8). Small changes in local composition, interaction strength, or connectivity to surrounding structures could therefore remain modest during interphase and become more apparent when the matrix disperses and reforms. We did not measure viscosity, surface tension, molecular exchange, or Rubisco–EPYC1 binding. A direct physical interpretation will require measurements such as fluorescence recovery after photobleaching or biochemical reconstitution.

Machine learning served as a discovery tool rather than as a replacement for biological inspection. We did not systematically benchmark the CAE-OC-SVM against hand-crafted morphology metrics and therefore do not infer that it outperforms feature-based analysis. Its practical role was to apply a common unsupervised scoring framework before mutant classes were known. Visual inspection remained important, as illustrated by *pim8*, which showed a modest anomaly rate but a reproducibly small pyrenoid.

The genetic results provide different levels of support. For most *pim* strains, insertion-site mapping only nominates candidate loci, and additional lesions in an insertional background cannot be excluded. *pim13* mapped to EPYC1, consistent with the established role of EPYC1 in matrix assembly. *pim8* contained three mapped insertions, one of which was in *STT7*. Six independent *STT7*-edited clones generated at two target sites yielded PCR amplicons consistent with target-site modifications and lacked detectable STT7 accumulation; three selected clones also showed differential reconstruction-error patterns in the pyrenoid region. These observations support an association between loss of STT7 and altered pyrenoid image features, but they do not establish that *STT7* is the sole causal lesion underlying *pim8* or that STT7 acts directly on the pyrenoid. STT7 is known for its role in LHCII phosphorylation and state transitions (19, 20), whereas the recently described kinase KEY1 more directly regulates pyrenoid condensate size and cell-cycle-dependent dissolution by modulating Rubisco–EPYC1 interactions (14). The present data do not place STT7 in the same mechanistic category as KEY1. They instead suggest that photosynthetic or thylakoid state may influence pyrenoid organization indirectly.

The growth phenotypes also require cautious interpretation. Some *pim* strains grew slowly under HC, whereas others differed most clearly as CO_2_ supply decreased. The collection therefore does not define a single phenotype specific to the lowest CO_2_ condition. Because the assays were performed on MOPS-P medium without an added organic carbon source, they measured photoautotrophic growth across the imposed CO_2_ regimes. Even so, reduced growth cannot be assigned specifically to impaired CCM function, because broader defects in photosynthesis, chloroplast physiology, metabolism, or cellular growth could produce similar outcomes. The spot assays nevertheless provide an independent phenotypic dimension for comparison with image and division phenotypes. Identifying genes required for pyrenoid-based CCM performance as CO_2_ becomes limiting remains relevant to efforts to understand and engineer such systems in plants, but direct measurements of inorganic-carbon affinity or photosynthetic performance would be needed before individual *pim* loci could be assigned such roles (21, 22).

Several additional limitations define the scope of the study. The known-mutant panel provided biological calibration but not an independent estimate of classifier sensitivity or specificity. The pooled screen and visual prescreen favored reproducible phenotypes and may have missed low-penetrance defects. Reconstruction-error maps localize differences in reconstruction performance but do not identify the responsible molecule, demonstrate gain or loss of a structure, or provide a statistical significance map. Fluorescence intensity can contribute to the learned representation, so an anomaly score cannot be interpreted as shape alone. The live-imaging analysis was qualitative and was not used to estimate genotype-level effect sizes or statistical differences. Finally, all comparisons used the same RBCS1-mGold reporter background, which controls for reporter effects within the study but does not exclude an influence of the reporter itself.

In summary, unsupervised image analysis expanded a forward genetic screen beyond phenotypes that could be specified in advance. The resulting *pim* collection includes strains with subtle interphase image phenotypes and diverse mitotic or growth phenotypes and provides candidate loci for further genetic analysis. Similar strategies may be useful for other dynamic cellular compartments when informative phenotypes are difficult to define in advance or emerge most clearly during remodeling.

## Materials and Methods

### Strains and pyrenoid reporter

The RG2 reporter strain was generated in a wild-type *Chlamydomonas reinhardtii* CC-125 background to express a Rubisco small subunit 1 (RBCS1)-mGold fusion protein for visualization of the pyrenoid matrix. The same reporter background was used for screening, differential reconstruction-error mapping, live imaging, and growth assays unless otherwise indicated. Established pyrenoid-related mutants, including *epyc1*, *saga1*, *saga2*, *mith1*, *rbmp1*, and *rbmp2*, were used for biological calibration of the anomaly-detection pipeline. Unless otherwise noted, cells were grown in Tris-acetate-phosphate (TAP) medium at 25°C under continuous illumination (∼120 μmol photons m^−2^ s^−1^) with gentle shaking (120 rpm). Once cultures reached mid-logarithmic phase (optical density at 730 nm [OD_730_] = 0.4–0.7), cells were harvested by centrifugation (2,000 g, 5 min) and transferred to MOPS-buffered phosphate (MOPS-P) medium for experimental treatments.

### Image preprocessing and dataset construction

Confocal laser-scanning microscopy images of RG2 cells were processed with StarDist2D to extract individual cell regions (18). Objects were excluded when they were within 10 pixels of the image boundary, had cell areas outside 200–8,000 pixels^2^, or had eccentricity ≥0.95. After filtering, 4,905 WT single-cell images were retained. Images were standardized by contrast-limited adaptive histogram equalization (clip_limit = 0.02) and resized to 64 × 64 pixels. Source image files were partitioned before augmentation, with 80% assigned to training and 20% to validation, so that augmented derivatives from one imaging field remained within the same partition. A 22-fold augmentation scheme used rotations within ±2°, translations within ±2%, and horizontal or vertical flipping, producing 107,910 image instances in total.

### Convolutional autoencoder and OC-SVM

The encoder contained three convolutional blocks with 32, 64, and 32 filters, 3 × 3 kernels, batch normalization, rectified linear unit activation, and 2 × 2 max pooling. It compressed each 64 × 64 image into a 2,048-dimensional latent vector. The decoder reconstructed the input through symmetric upsampling and convolutional layers. The model was trained with mean squared reconstruction loss using the Adam optimizer at a learning rate of 0.001. Early stopping with a patience of 10 epochs was based on validation loss; training stopped after 22 epochs.

Latent feature vectors were standardized and reduced from 2,048 to 100 dimensions by principal component analysis. A one-class support vector machine with a radial basis function kernel was fit to the WT features (17). The parameter ν was set to 0.05. For each strain, the anomaly rate was the fraction of cells classified outside the estimated WT support. Relative anomaly rates were calculated by dividing the anomaly rate of each strain by the WT anomaly rate when used for figure display.

### Biological calibration, hybrid mutant screening

Established pyrenoid-related mutants were processed using the same segmentation, preprocessing, encoding, dimensionality-reduction, and classification workflow applied to WT and *pim* strains. This panel was used to determine whether established pyrenoid phenotypes produced biologically interpretable anomaly rates. It was not treated as an independent test set for estimating classifier sensitivity or specificity.

For the mutant screen, approximately 21,000 random insertional mutants were visually surveyed to exclude strains with obvious growth defects or complete loss of the pyrenoid reporter. The remaining 1,728 strains were pooled at 24 colonies per well, producing 72 pools. Pools whose anomaly rates exceeded the matched WT baseline by approximately 4.2 percentage points were selected for secondary single-colony analysis. Individual colonies from selected pools were re-imaged and retained when their anomaly rates reproducibly exceeded 1.5 times the WT baseline or when repeated visual inspection revealed a consistent pyrenoid phenotype. WT controls were included across screening batches to monitor day-to-day and plate-to-plate variation.

### Differential reconstruction-error mapping

For reconstruction-error mapping, WT source image files were divided into an 80% reference subset and a separate 20% comparison subset. Unaugmented images were used. For each cell, the raw reconstruction-error map was calculated as the absolute pixel-wise difference between the model-input image and its CAE reconstruction. The mean raw reconstruction-error map of the 80% WT reference subset was used as the WT reference map. A differential reconstruction-error map was generated by subtracting this reference map from the raw map of each cell. The mean WT differential map shown in Fig. 3C was calculated by averaging the differential maps of the 20% WT comparison subset. For genotype-level displays, aligned differential maps were averaged across cells of the indicated genotype. The corresponding aligned chlorophyll-autofluorescence and RBCS1-mGold images were averaged separately and merged to provide morphological context. These mean fluorescence images were not used for quantitative comparisons of fluorescence intensity.

### Four-dimensional live imaging

Cells were immobilized in low-melting-point agarose and imaged by confocal laser-scanning microscopy. Time-lapse image stacks of RBCS1-mGold fluorescence were used to follow matrix dispersal, formation and distribution of Rubisco-containing foci, and post-division pyrenoid recondensation.

Imaging was performed using a Leica SP8 confocal microscope, with a 63× objective. mGold was excited at 514 nm, and fluorescence emission was collected at 520–540 nm. Z-stacks were acquired at 0.3 µm intervals (20 optical sections per stack), with a time interval of 60 sec between time points. Imaging was performed at room temperature, with a total observation period of <300 min. Representative time-lapse series are provided as Supplementary Movies.

### CRISPR-Cas9 system-mediated generation of *STT7* mutants

To generate *stt7* mutants, we designed two guide RNAs (5′-CTCCTGTAGACCGCTCAATG-3′ and 5′-GGTGCTGCAATCTGCTCCAG-3′) targeting the first exon of the *STT7* gene using CRISPOR (http://crispor.tefor.net/). The crRNA and tracrRNA were chemically synthesized (Integrated DNA Technologies) and mixed with recombinant Cas9 protein to form a ribonucleoprotein (RNP) complex. RNP and the *AphVII* cassette (conferring hygromycin resistance) were co-introduced into the RG2 reporter strain by electroporation (23).

Transformants were selected on TAP plates containing 15 µg mL⁻¹ hygromycin and screened by colony PCR. Specific genomic insertion of the a*phVII* cassette was verified by PCR with primers 5′-CGTGTAAGGAATAGAACGTGTATTGTATGC-3′ (F1) and 5′-CTGCACGAGAAATGACCAACACAGATC-3′ (R1). Immunoblotting with a STT7-specific antibody (Agrisera) confirmed the loss of STT7 protein in the resulting mutant. Histone H3 was used as a loading control.

### Insertion-site mapping

Genomic DNA samples were submitted to Novogene for plant whole-genome sequencing. Whole-genome libraries were prepared using the provider’s standard non-PCR-free library preparation workflow and sequenced on a NovaSeq X Plus instrument to generate 150-bp paired-end reads, targeting 6 Gb of sequence data per sample. Samples were submitted under the provider’s stated quality requirements of at least 100 ng genomic DNA in a volume of at least 20 µL, at a concentration of at least 5 ng/µL, with an OD_260_/OD_280_ ratio of 1.8–2.0 and no detectable degradation or contamination.

Sequencing reads were analyzed against the *Chlamydomonas reinhardtii* reference genome assembly v5.6. Insertion sites were identified from insertion-supporting reads and assigned relative to annotated gene models. Because complementation, segregation analysis, or independent alleles were not available for most strains, mapped loci were treated as candidate loci rather than causal genes. Mapped insertion sites and associated candidate loci are listed in Table 1.

## Data, code, and materials availability

Custom scripts used for the machine-learning analyses are publicly available at https://github.com/Yamano-Lab/2025_Machine_Learning-based_screening. Other data supporting the findings of this study, including anomaly scores, source measurements, raw and processed microscopy files, uncropped immunoblots, and insertion-site mapping records, are available from the corresponding author upon reasonable request. Strains and other materials generated in this study are available from the corresponding author subject to applicable institutional agreements.

## Acknowledgments

We thank Takenobu Ogawa for insightful discussions. We thank Ryutaro Tokutsu, Jerome Gunasekran, Nahoko Nishimura, Kazuko Tarui, Hatsue Mizuhara, and Hiroko Shimada for their technical assistance in conducting the experiments.

